# First report of marine sponge *Chelonaplysilla delicata* (Demospongiae: Darwinellidae) from the Andaman Sea/Indian Ocean with a baseline information of epifauna on a mesophotic shipwreck

**DOI:** 10.1101/636043

**Authors:** Rocktim Ramen Das, Titus Immanuel, Raj Kiran Lakra, Karan baath, Ganesh Thiruchitrambalam

**Affiliations:** Department of Ocean Studies and Marine Biology, Pondicherry University, brookshabad, Andaman Islands, India; Graduate School of Engineering and Science, University of the Ryukyus, Nishihara, Okinawa, Japan; Marine Biology Regional Centre (MBRC), Zoological Survey of India (ZSI), Chennai, Tamil Nadu, India; Infinity Scuba Andaman’s, Chidiyatapu, Andaman Islands, India

**Keywords:** Biodiversity, Porifera Taxonomy, Invasive, *Tubastraea*

## Abstract

During a biodiversity assessment on a wreck located in the Andaman Sea (Andaman Islands), a single specimen of sponge *Chelonaplysilla delicata* was recorded. Our finding confirms the species taxonomy and highlights the current observation as a first report from the Andaman Sea/ Indian Ocean. The baseline information of epifauna is further stated in this study.

## INTRODUCTION

The Andaman Sea, an eastern subdivision of the Indian Ocean is bordered by countries like Thailand, Myanmar on the east and the Andaman archipelago (Andaman & Nicobar Islands/ANI) on the west (Brown 2007). (Figure 1). A large portion although falls within the boundary of the Coral Triangle Initiative (CTI) (Rudi et al. 2012) studies related to its marine biodiversity or the coral reef ecosystem has been comparatively understudied or scattered (Aungtonya et al. 2000; Brown 2007). Additionally, the Andaman Sea possesses several shipwrecks (Kheawwongjan & Kim 2012) acting as artificial reef ecosystems knowledge pertaining to which is mostly limited in the region. These sunken structures provide space for growth and establishment of various sessile marine communities like poriferans (Walker et al. 2007; Lira et al. 2010) and other non-native species (Patro et al. 2015; Soares et al. 2020). Within the Indian EEZ, recent studies targeting shallow-water wrecks has filled important knowledge gaps (example Mohan 2013; Das 2014; Yogesh-Kumar et al. 2015) (Table 1). This article further adds essential information in these rarely studied ecosystems at a mesophotic depth and reports a marine sponge from the Andaman Sea/Indian Ocean.

**Figure 1.**
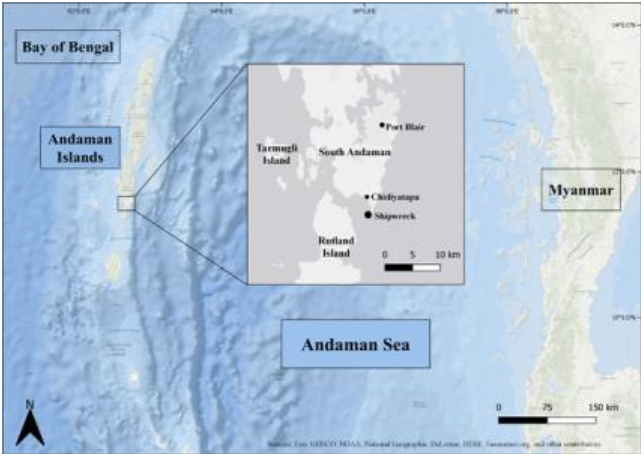
Location of the study area

**Table 1.** Information on biodiversity studies in a wreck ecosystem in Andaman and Nicobar Islands

| Wreck Name | Co-ordinates | Location | Date of Sinking | Depth (m) | Current Activities | Reference |
| --- | --- | --- | --- | --- | --- | --- |
| SS Inchkeith | 12°00'23.69"N<br>92°46'08.34"E | Kyd Island (South Andaman) | 1955 | 14 | Scuba | Mohan 2013 |
| **HMIS Sophie Marie** | **11°28'38.02"N**<br>**92°42'12.20" E** | **Chidiyatapu (South Andaman)** | **1942** | **30 - 33** | **Scuba** | **Present Study** |
| MV Mars | 11°55'54.98"N<br>92°57'24.12"E | Havelock (Ritchie's Archipelago) | 2006 | 10 - 16 | Scuba | Das R. R. (Pers. Obs.) |
| North Bay Wreck | 11°43'00.56"N<br>92°45'60.60" E | Port Blair (South Andaman) | 30 – 40 (yrs) | 10 | Scuba and Fishing | Yogesh-Kumar et al. 2015 |
| Peel Wreck | 12°03'84.20"N<br>92°57'81.10" E | Havelock (Ritchie's Archipelago) | 8 - 10 | 9 - 12. | Scuba | Yogesh-Kumar et al. 2015 |
| Japan Wreck | 09°10'88.30"N<br>92°50'12.30" E | Car Nicobar (Nicobar Islands) | 40 - 50 | 28 | Fishing | Yogesh-Kumar et al. 2015 |
| Sinclair Bay Shipwreck | 11°39'873"N<br>92°45'488"E | Near Ross Island (South Andaman) | - | 8 |  | Mondal & Raghunathan 2017 |

## MATERIAL AND METHODS

The sponge *C*.*delicata* (Image 1), collected from the shipwreck HMIS SM* during a survey conducted for documenting epifaunal diversity from February to March 2014. The wreck is a 70m long Royal Indian navy minesweeper that sank in the year of 1942. At a depth of ≈33 meters, it lies at the edge of the Macpherson strait near Chidiyatapu (11°28’38.02” N 92°42’12.20” E) (Figure 1). Water transparency and temperature were recorded with Secchi disc and dive calculator. Within 2 hours after collection, the specimen was preserved in 100% ethanol. A surface peel of the easily separable cortex of the specimen was removed and placed in xylene for 24 hours after which a permanent slide of the peel was mounted with DPX. A single fibre with its base and branches intact was removed from the sponge for species-level identification under a stereo microscope. (Image 1B-D). The specimen was identified following Pulitzer-Finali & Pronzato (1999). The preserved specimen is deposited in the National Zoological Collections (NZC) of the Andaman and Nicobar Regional Centre (ANRC), Zoological Survey of India (ZSI), Port Blair.

Benthic cover was assessed by randomly placing 20 (0.25×0.25cm) quadrats (Image 2). The photographs were analysed using open-sourced Coral-Net software (Beijbom et al. 2012) and the epifauna was categorized into Unknown, Sponge, Hard Coral, *Ircinia* sp., Algae, *Iotrochota* sp., Sediment, *Tubastraea* aff. *coccinea, Tubastraea micranthus*, Hard Substrate, Ascidian, Bleached Coral (modified from Zintzen et al. 2006). Data from the annotated quadrats was later transferred and processed in Microsoft Excel^®^ (Microsoft 365 MSO, 16.0.13001.20266/32bit). Study maps were created using open-sourced Quantum Geographic Information System (QGIS ver. 3.6).

## RESULTS AND DISCUSSION

*Systematics*

Phylum: Porifera

Class: Demospongiae

Order: Dendroceratida

Family: Darwinellidae

Genus: *Chelonaplysilla*

Species: *C. delicata* Pulitzer-Finali & Pronzato, 1999

Paratype: 1 ex (Paratype)., ZSI/ANRC – 14321, India: Andaman Island: South Andaman: Chidiyatapu (11°28’38.02” N; 92°42’12.20” E). Coll. Rocktim Ramen Das, 2014

### Diagnosis

*C*.*delicata* predominantly thickly encrusting (< 10 mm) but has erect lobes that are about 4 - 5 cm high. The sponge surface is conulose, and the acute conules separated from each other by 2 - 5 mm. Oscules 1 - 3 mm in diameter, flush with the surface and unevenly distributed all over on sponge surface. The texture is soft collapsible and feeble. The fresh specimen was dark violet or purple in colour and retained its colour even in the preserved condition. Sponge surface covered by structured regular reticulation of sand and spicule detritus, which forms regular roundish or oval meshes of 90 - 155 µm. This reticulation is typical of the genus. Regular rounded fibrous pores, inhalant in nature, is enclosed within these rounded meshes (Image. 1D). The skeleton is dendritic, made up of pigmented fibres fragile in nature with repeated branching that originate from a basal spongin plate (Image 1B-C) and extends towards the boundary. The primary fibre measured to be around 0.4 mm at its thickest. Spicules are absent.

#### Distribution

India: Andaman Sea (ANI, South Andaman, Present study). Elsewhere: Bismark Sea (Papua New Guinea) (Pulitzer-Finali and Pronzato, 1999), Indonesia (Sulawesi) (GBIF, 2000), Palau (Micronesia) (Ridley et al. 2005), French Polynesia (Alencar et al. 2017) (Figure 2).

**Figure 2.**
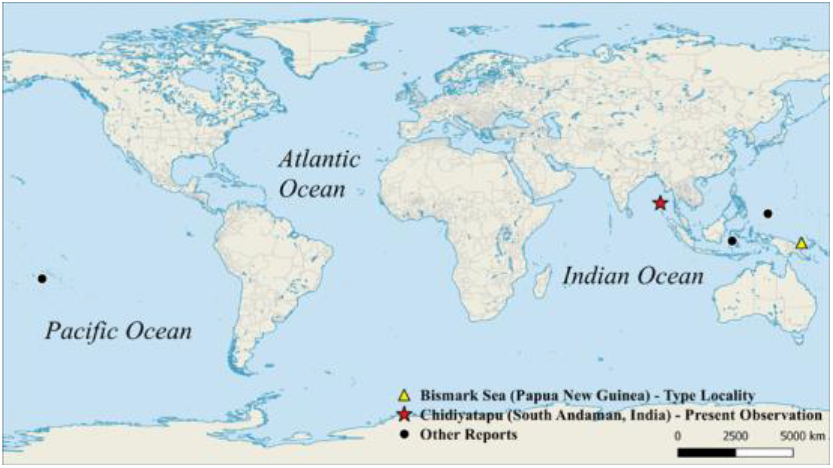
Global distribution of *C. delicata* Pulitzer-Finali & Pronzato, 1999

**Figure 3.**
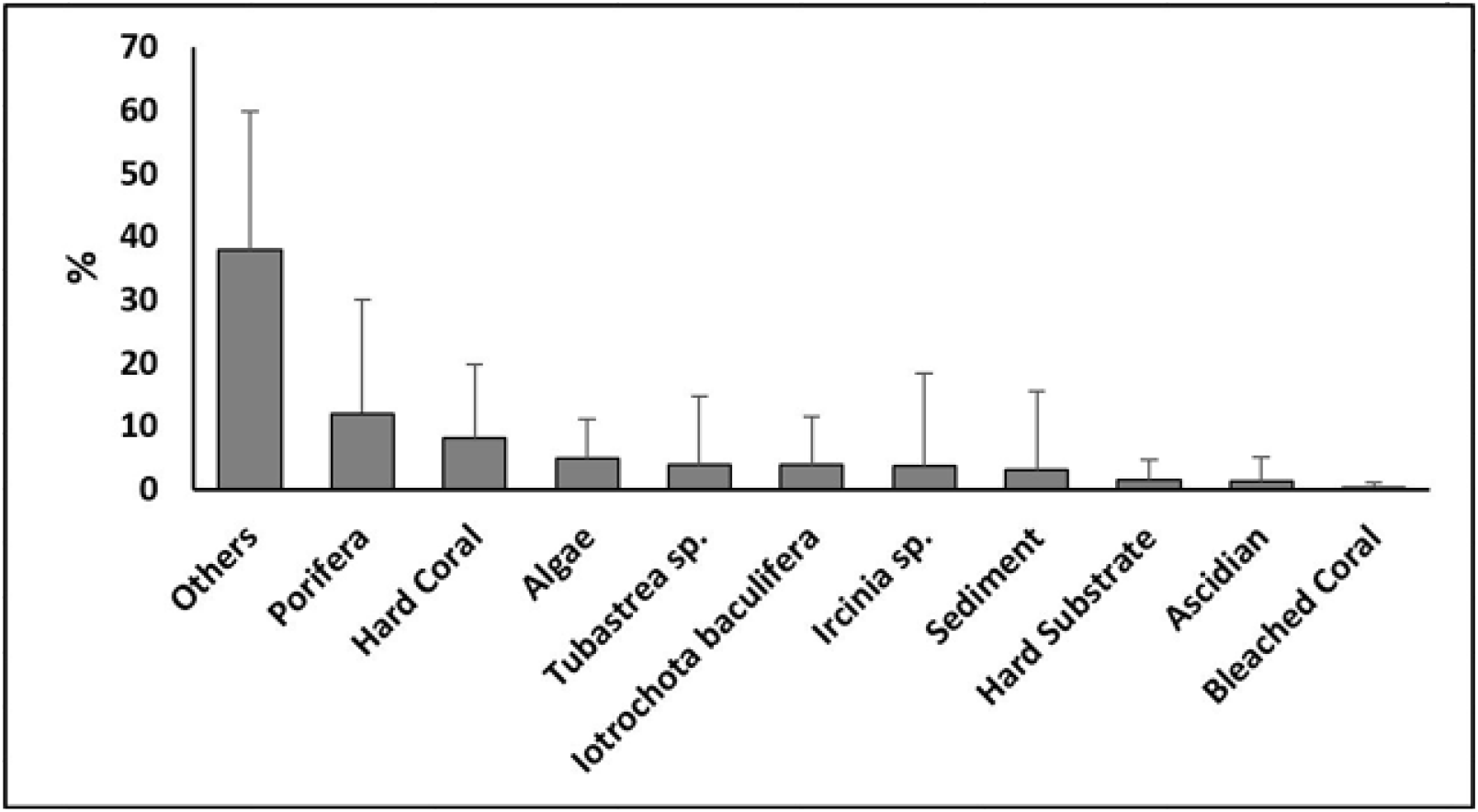
Mean percent cover of epifauna obtained from twenty 0.25×0.25 quadrats

#### Remarks

*C*.*delicata* is very similar to *C*.*erecta* (Tsurnamal, 1967); however, the latter has fibres anastomosing in nature whereas the thickness of fibres in *C*.*delicata* fades in diameter. The specimen mentioned in Pulitzer-Finali and Pronzato, 1999 is Gray whereas our specimen in dark maroon in live condition (Image 1A). The specimen was initially misidentified as *C*.*erecta* (Das 2014; Das et al. 2016), thus there was a need for the update and fill the knowledge gap of this species distribution range.

#### Comments

The family Darwinellidae possesses sponging fibres with proper skeleton and fibrous spicules (Van Soest 1978; Bergquist & Cook 2002). It consists of four recognized genus and forty-seven accepted species (one under “nomen nudum” status). *Chelonaplysilla* is the only genus, which is devoid of spicules but consists of a fibrous dendritic skeleton that possesses distinct laminated bark surrounding a central pith region. A structured and separable cortex that is reinforced by a delicate reticulation of sand grains (Van Soest 1978) distinguishes this genus.

#### Wreck Biodiversity

Benthic cover assessment (Image 2) reveal that sponges are the second most abundant group on the surface of the wreck mostly encrusting in nature. In most instances the encrusting sponge genus *Iotrochota* was readily visible. Ahermatypic and invasive sun corals were abundant in selected localities and may have found a successful substrate for further expansion (Image 2A, 3). Few polyps of *Tubastraea micranthus* had bleached, a stark contrast to their ahermatypic nature. Updated and revised identification following Das et al. (2016) on the wreck surface includes sclectenian genus *Favia, Symphyllia, Podabacia crustacea*, and *Leptoseris*. A single individual of the Gastropod genus (*Chicoreus*), and few Crinoids. The identified Poriferan families include Irciniidae (*Ircinia*), Chalinidae (*Haliclona* (*Reniera*)); Thorectidae (*Hyrtios*), Iotrochotidae (*Iotrochota baculifera*), Thorectidae (*Dactylospongia*). Tunicates comprised of Didemnidae (*Didemnum*), Perophoridae (*Perophora*) and other unidentified sp.

The Faunal organisms that thrive in Artificial Reefs (Shipwrecks) is an important part of the marine community (Amaral et al. 2010; Zintzen et al. 2006). With increasing anthropogenic impacts on natural coral reef habitats, artificial reefs are regarded as a successful alternative (Perkol-Finkel & Benayahu 2005). As a result, it becomes important to understand the biological communities growing on these habitats (Thanner et al. 2006). Sponges which naturally occupy shipwrecks are one of the dominant organisms in such habitat, as evidenced in the present study, however its diversity will be strictly limited to the environmental settings. For example, some species of genus *Iotrochota* is found in sheltered environments (Cleary & Voogd 2007) as seen in our observation (Image 2C). Similarly, shipwrecks are also known to act as a successful substrate for many non-native species as reported from the Atlantic and the Red Sea (Perkol-Finkel et al. 2006; Soares et al. 2020). Repeated encounter of *Tubastrea* aff. *coccinea* and T. *micranthus* in the study site is a strong evidence from the Andaman Sea (Figure 1). The sponge species reported in herein is in a much-extended depth compared to its initial described type locality (see. Pulitzer-Finali & Pronzato 1999).

Technical difficulties are a foremost reason for these habitats to have limited studies at mesophotic depths (Massin et al. 2002; Zintzen et al. 2006), but with the rapid scale development of ROV’s and submersibles, detailed exploration of these ecosystems can be well predicted. Further these areas might be a hub for various underexplored flora and fauna and might be effective in reviving threatened marine life due to the loss of natural ecosystems.

## Author Contribution Statement

### Rocktim Ramen Das

Conceptualization, Study Design, Writing – Original Draft, Reviewing & Editing, Field Assessment, Sample Collection, Data Analysis, Laboratory Analysis.

### Titus Immanuel

Writing Original Draft, Reviewing & Editing, Sample Identification, laboratory Analysis.

### Raj Kiran Lakra

Study Design; Writing, Reviewing & Editing, Field Assessment.

### Karan Baath

Field Assessment

### Ganesh Thiruchitrambalam

Study Design; Writing, Reviewing & Editing; Supervision.

## Acknowledgements

The authors thank Dr. P.M. Mohan, Department of Ocean Studies and Marine Biology (DOSMB), Pondicherry University, India for the necessary facilities. The authors acknowledge Dr. S.Venu (DOSMB, Pondicherry University, India) for initial comments on the project. Drs. E. Hirose and F. Sinniger (University of the Ryukyus, Japan) for valuable comments regarding this draft. Drs. K. Wangkulangkul (PSU, Thailand), S.Y. Tenjing (NCSCM, India) for English editing.

---

**Image 1.**
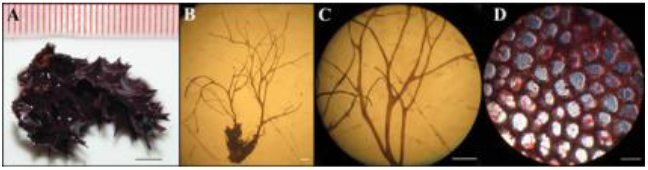
*Chelonaplysilla delicata* [ZSI/ANRC-14321]: (A) Freshly collected specimen, (B) Branching fibres and basal sponging plate, (C) Closer view of pigmented, branching, dendritic spongin fibre, (D) Inhalant pores surrounded by rounded meshes reinforced by sand grains. Scale (A) 5mm (B) 2 mm, (C) 2 mm, (D) 155 μm.

**Image 2.**
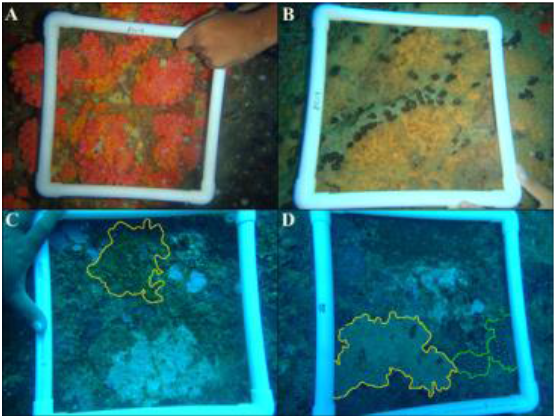
A) *Tubastraea* aff. *coccinea* B) *Ircinia* sp. C) Mixed assemblage of communities. Encrusting sponge *Iotrochota* sp. (Yellow) D) Mixed assemblage, Sponge (Yellow), coral (green).

**Image 3.**
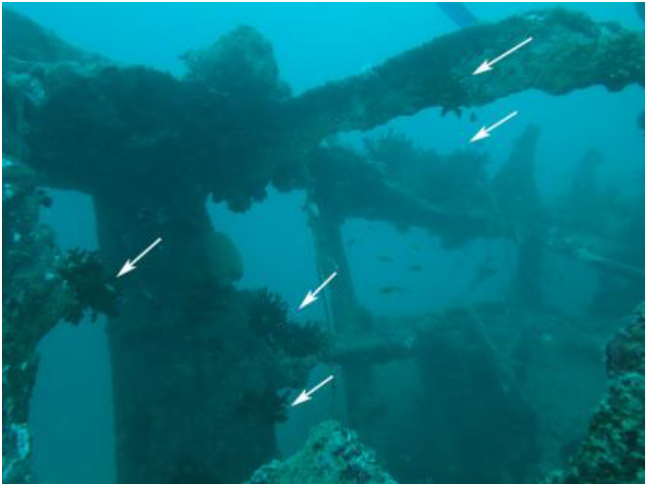
A part of the wreck HMIS SM. (Arrow: high abundance of invasive *Tubastraea micranthus*)

